# RBM15 promotes lung adenocarcinoma progression and palbociclib sensitivity *via* m6A-mediated STIL activation and downstream cyclin D1/CDK4 signaling

**DOI:** 10.64898/2025.12.04.692261

**Authors:** Yifan Chen, Jiaojiao Zhang, Hongjuan Yue, Lihong Wang, Minghong Lv, Ling Wu, Junling Ma, Jing Wang, Weifang Yu, Ting Liu, Jia Wang

## Abstract

RNA-binding motif protein 15 (RBM15), an N6-methyladenosine (m6A) methyltransferase, plays a crucial role in the progression of lung adenocarcinoma (LUAD); however, the underlying mechanisms remain insufficiently defined. In this study, we investigated the contribution of RBM15-mediated m6A modification to LUAD pathogenesis. Our data showed that RBM15 was highly expressed in LUAD tissues and cell lines. Notably, its expression levels were positively correlated with tumor TNM staging and negatively correlated with differentiation status. The overexpression of RBM15 enhanced cell proliferation, migration, invasion, and cell cycle progression, while simultaneously inhibiting apoptosis. Conversely, RBM15 knockdown reversed these phenotypes and induced G2/M-phase arrest. *In vivo* experiments further confirmed that RBM15 knockdown led to the suppression of tumor growth. RNA sequencing, MeRIP sequencing, and bioinformatics analyses identified *STIL* as a gene directly regulated by RBM15 through m6A modification, a finding subsequently validated by luciferase reporter assays. Western blot and rescue experiments demonstrated that RBM15 and STIL cooperatively activate the cyclin D1/CDK4 signaling pathway, thereby promoting LUAD progression. However, this effect was markedly attenuated by CDK4/6 inhibitor application. Collectively, these findings indicated that the RBM15-STIL-cyclin D1/CDK4 axis represents a promising therapeutic target for LUAD.

## Introduction

Lung cancer is among the most aggressive malignancies and remains the leading cause of cancer-related mortality worldwide. According to the National Cancer Report of 2022, approximately 828,000 new lung cancer cases were recorded in China that year, resulting in 657,000 deaths. These figures represent 20% of all cancer incidences and 27% of all cancer-related mortality[2], respectively, thus ranking LUAD first in both categories. Lung cancer is often asymptomatic in its early stages, leading to delayed diagnosis and, consequently, poor prognosis, with an overall 5-year survival rate of only 16.8%[3]. Pathologically, the disease is broadly classified into small-cell lung cancer (SCLC) and non-small cell lung cancer (NSCLC). Among the NSCLC subtypes—adenocarcinoma, squamous cell carcinoma, and large cell carcinoma—lung adenocarcinoma (LUAD) is the most prevalent, accounting for approximately 40% of all lung cancer cases. The increasing incidence of LUAD is partly attributed to cigarette smoking, environmental nitrate exposure, and a rising prevalence among women. Its high mortality rate is associated with difficulties in early detection, the potential for rapid metastasis, resistance to therapy, and the lack of effective systemic treatments[1]. Although the etiology and pathogenesis of LUAD remain incompletely understood, growing evidence indicates that it arises from the interplay of multiple factors, including smoking, environmental pollutants, genetic susceptibility, and exposure to carcinogenic compounds such as polycyclic aromatic hydrocarbons and heavy metals. Over recent years, epigenetic regulation, which significantly influences gene expression without altering the DNA sequence, has also been established as a crucial contributor to LUAD initiation and progression. Epigenetic mechanisms, and, in particular, RNA modifications such as the addition of N6-methyladenosine (m6A), have been increasingly recognized as key modulators of tumor biology. Therefore, elucidating the molecular and epigenetic mechanisms underlying LUAD pathogenesis is essential for identifying novel diagnostic biomarkers and therapeutic targets for this malignancy, as well as for advancing precision medicine in lung cancer.

M6A is the most prevalent internal RNA modification, regulating gene expression by modulating RNA stability, splicing, and translation. This modification governs diverse biological processes, including cell differentiation, development, and tumorigenesis[4]. Its deposition and removal are dynamically controlled by a complex regulatory network consisting of methyltransferases (“writers”), demethylases (“erasers”), and binding proteins (“readers”). RNA-binding motif protein 15 (RBM15), a core component of the m6A methyltransferase complex, interacts with METTL3, METTL14, and WTAP to catalyze m6A installation[5]. Emerging evidence has implicated RBM15 as an oncogenic driver in multiple malignancies. For example, RBM15 was found to promote laryngeal squamous cell carcinoma progression through IGF2BP3-mediated m6A site recognition[6], as well as enhance hepatocellular carcinoma growth *via* IGF2BP1[7]. Consistent with these observations, its depletion was reported to suppress colorectal cancer proliferation and metastasis[8]. However, the role of RBM15 in LUAD remains unclear.

In this study, we found that RBM15 expression is significantly elevated in LUAD, with high expression correlating with poor prognosis. Functional analyses further revealed that RBM15 promotes LUAD cell proliferation and metastasis both *in vitro* and *in vivo*. Mechanistically, we identified the SCL-interrupting locus (STIL) protein as a previously unrecognized downstream effector of RBM15. Acting in concert, RBM15 and STIL drive LUAD progression *via* the activation of the cyclin D1/CDK4 signaling pathway. Notably, the CDK4/6 inhibitor palbociclib effectively reversed these effects, highlighting the therapeutic relevance of the RBM15-STIL axis. Collectively, our findings define RBM15 as a potential oncogenic regulator and therapeutic target in LUAD.

## Results

### The expression of RBM15 is significantly upregulated in LUAD tissues and is associated with poorer prognosis

Analysis of TCGA data revealed that the mRNA levels of RBM15 were significantly higher in LUAD tissues than in adjacent normal tissues (Fig. 1A), a finding consistent with the pan-cancer analysis (Fig. 1B) and validation results from the GEO dataset GSE43458 (Fig. 1C). UALCAN analysis further demonstrated that RBM15 protein levels were also elevated in LUAD (Fig. 1D). Survival analysis using GEPIA2 indicated that high RBM15 expression was associated with significantly poorer overall survival (Fig. 1E). Given the limited number of studies on RBM15 in LUAD, these results prompted further exploration of its functional role in this malignancy. Immunohistochemistry results showed that RBM15 was primarily localized to the nucleus and cytoplasm, and its expression levels were significantly higher in LUAD tissues than in matched adjacent normal tissues (Fig. 1F, Supplementary Table 2). Clinicopathological analysis revealed that RBM15 expression was positively correlated with TNM staging and negatively correlated with differentiation grade, but was not associated with age, gender, tumor size, or lymph node metastasis. Consistent with these observations, qRT-PCR assay confirmed that RBM15 expression levels were higher in LUAD cell lines than in the BEAS-2B cell line. Based on their relative endogenous RBM15 expression levels, we selected A549 and H1975 cells for overexpression experiments, while H1299 and H1975 cells were used for knockdown experiments (Supplementary Fig. 1A). In summary, these findings indicated that RBM15 is upregulated in LUAD and is associated with aggressive clinical-pathological features, leading to poor prognosis.

**Fig 1.**
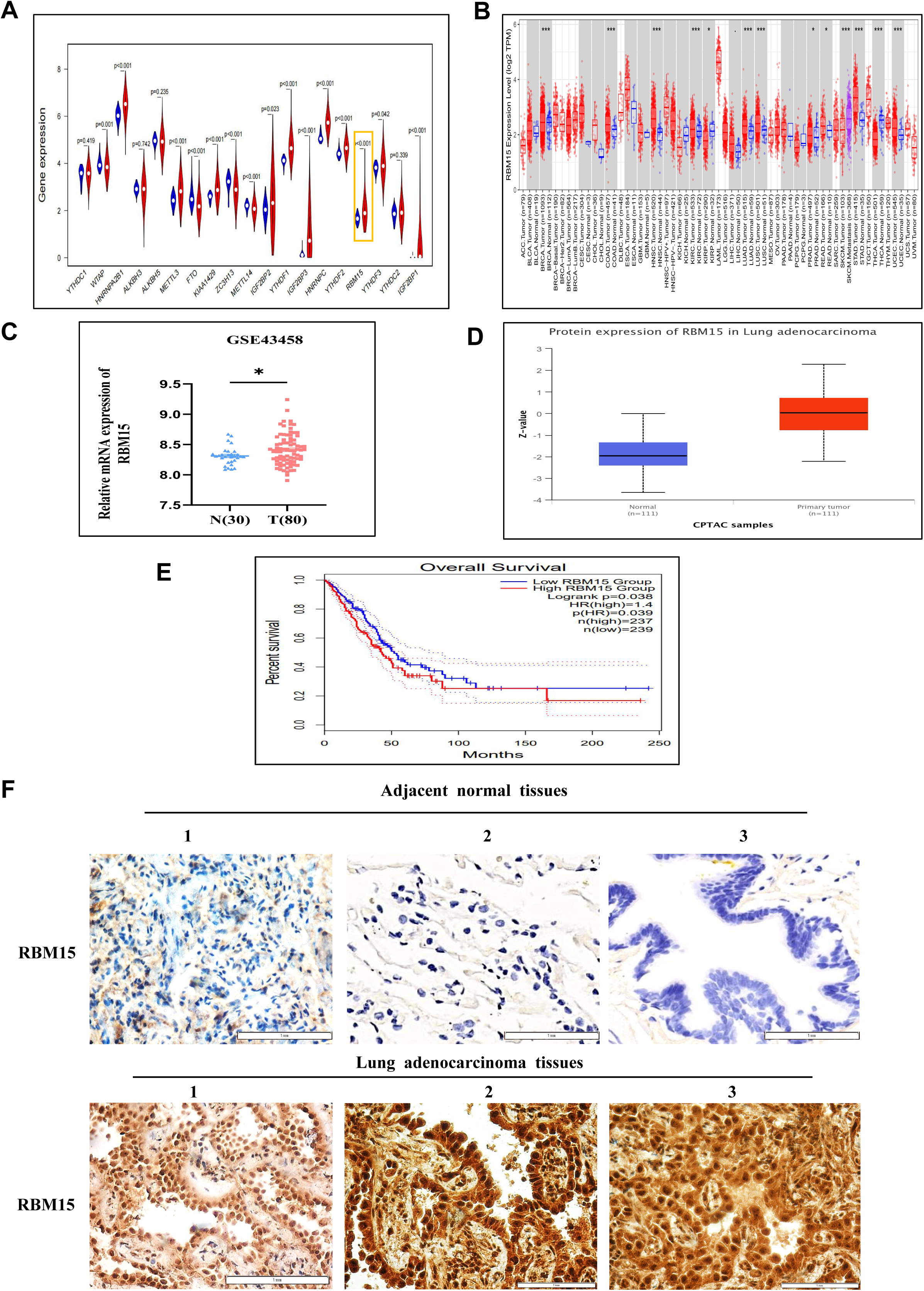
RBM15 is highly expressed in LUAD tissues and is associated with poor prognosis in patients with LUAD. (A) The expression levels of RBM15 in LUAD tissues and adjacent normal tissues based on TCGA data. (B) The expression levels of RBM15 in pan-cancer tissues and matched adjacent normal tissues based on data from TCGA database. (C) The expression levels of RBM15 in normal lung tissues and LUAD tissues from the GEO dataset GSE43458. (D) The protein expression of RBM15 in LUAD tissues and normal tissues based on the UALCAN database. (E) Survival curve analysis for RBM15 based on the GEPIA2 database. (F) Immunohistochemical detection of RBM15 expression (× 400 magnification). \**P* < 0.05, \*\*\**P* < 0.001.

### The overexpression of RBM15 promotes LUAD cell proliferation, migration, invasion, and cell cycle progression, while simultaneously inhibiting apoptosis

To determine the functional role of RBM15 in LUAD, we transfected A549 and H1975 cells with pcDNA3.1-RBM15 or the control vector. Successful overexpression was confirmed by qRT-PCR and western blot (Supplementary Fig. 1B, C). Cell viability (CCK-8) assays showed that RBM15 overexpression significantly enhanced cell proliferation compared with the control treatment (Fig. 2A, B). Correspondingly, wound healing and Transwell assays demonstrated that RBM15 markedly increased the migratory and invasive capacities of LUAD cells (Fig. 2C–F). Flow cytometry revealed that RBM15 overexpression reduced the rate of apoptosis and induced cell cycle arrest at the G1/S phase, as evidenced by an increase in the proportion of G0/G1 cells and a corresponding reduction in the fraction of S-phase cells (Fig. 2G, H). Collectively, these findings indicated that RBM15 acts as an oncogenic factor, promoting the growth, migration, and invasion of LUAD cells, while concurrently suppressing their apoptosis.

**Fig 2.**
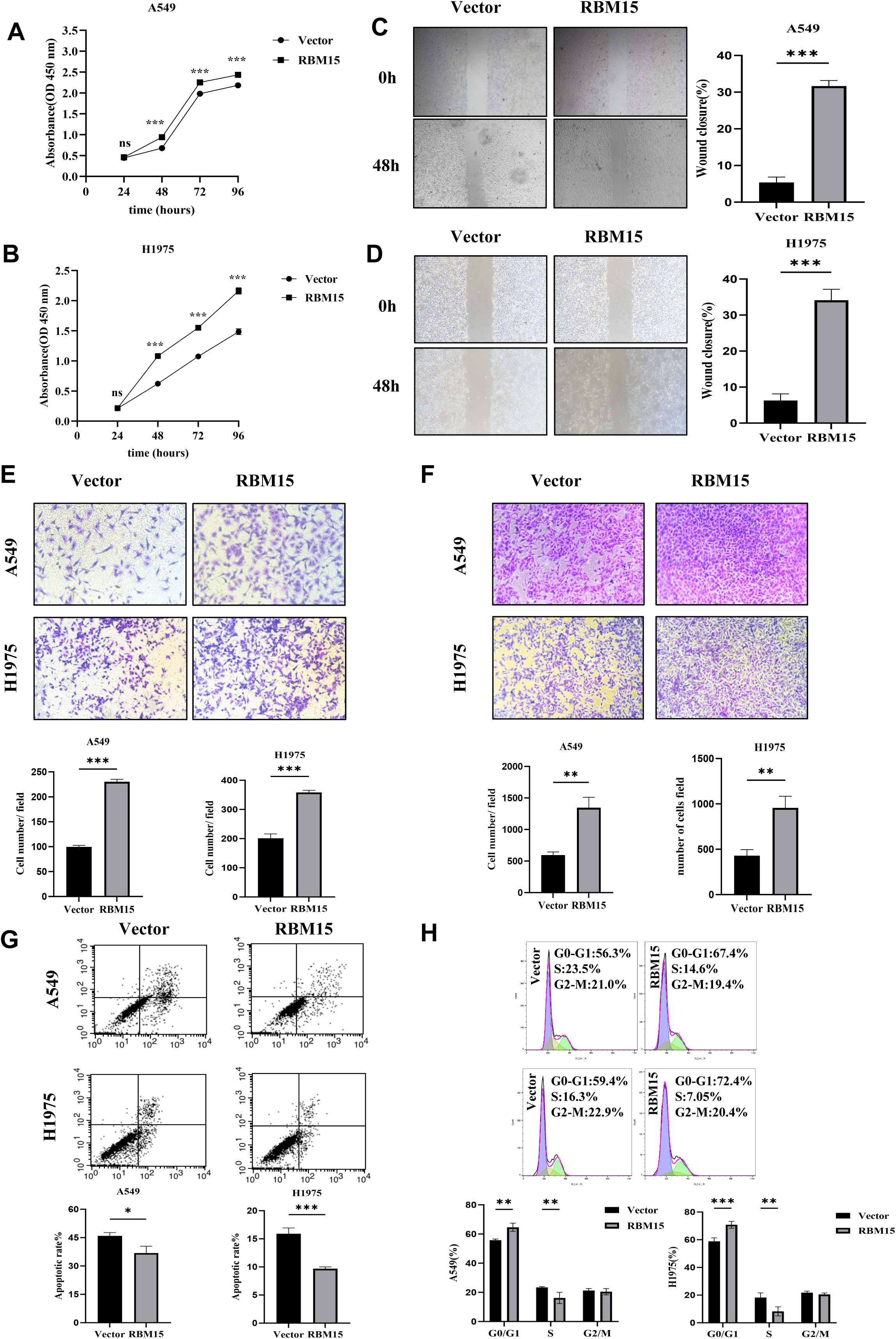
The overexpression of RBM15 promotes the proliferation, migration, and invasion of LUAD cells, inhibits their apoptosis, and induces cell-cycle arrest at the G1/S phase. (A, B) CCK-8 assays assessing the viability of A549 and H1975 cells. (C, D) Wound healing assays evaluating the migratory capacity of A549 and H1975 cells (× 40 magnification). (E, F) Transwell migration and invasion assays in A549 and H1975 cells (×100 magnification). (G) Flow cytometric analysis of apoptosis in A549 and H1975 cells. (H) Flow cytometric analysis of the cell cycle distribution in A549 and H1975 cells. \**P* < 0.05, \*\**P* < 0.01, \*\*\**P* < 0.001.

### RBM15 knockdown suppresses the proliferative, migratory, and invasive capacities of LUAD cells, promotes their apoptosis, and induces G2/M cell cycle arrest

To further validate the oncogenic role of RBM15 in LUAD, we silenced its expression in H1299 and H1975 cells using four siRNAs (si-1910, si-2116, si-2421, and si-2712). RBM15 silencing efficiency was confirmed by qRT-PCR and western blot (Supplementary Fig. 1D). CCK-8 assays revealed that RBM15 knockdown significantly inhibited LUAD cell proliferation (Fig. 3A, B). Wound healing and Transwell assays further demonstrated that RBM15 depletion markedly decreased LUAD cell migration and invasion (Fig. 3C–F). Flow cytometric analysis indicated that RBM15 silencing increased the rate of apoptosis in LUAD cells (Fig. 3G) and led to cell cycle arrest at the G2/M phase, as evidenced by a reduction in the proportion of G0/G1 cells and the accumulation of cells in the G2/M phase (Fig. 3H). These results confirmed that RBM15 knockdown suppresses the growth and metastatic potential of LUAD cells by promoting apoptosis and inducing G2/M phase arrest.

**Fig 3.**
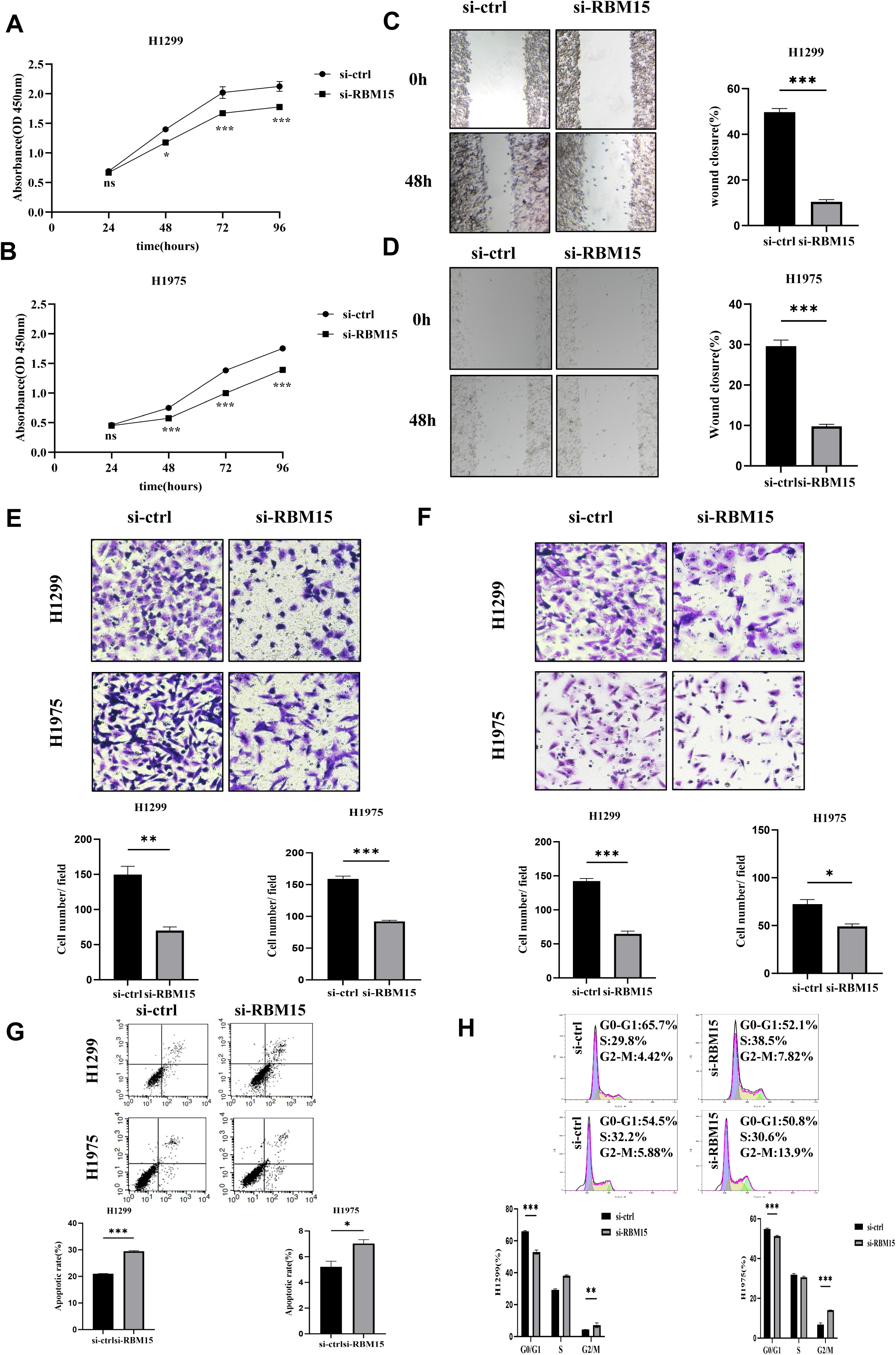
RBM15 knockdown inhibits the proliferation, migration, and invasion of LUAD cells, promotes their apoptosis, and induces cell-cycle arrest at the G2/M phase. (A, B) CCK-8 assays assessing the viability of H1299 and H1975 cells. (C, D) Wound healing assays evaluating the migratory ability of H1299 and H1975 cells (× 100 magnification). (E, F) Transwell migration and invasion assays in H1299 and H1975 cells (× 200 magnification). (G) Flow cytometric analysis of the apoptotic capacity of H1299 and H1975 cells. (H) Flow cytometric analysis of the cell cycle distribution in H1299 and H1975 cells. \**P* < 0.05, \*\**P* < 0.01, \*\*\**P* < 0.001.

### RBM15 depletion inhibits LUAD tumor growth *in vivo*

To examine the oncogenic potential of RBM15 *in vivo*, we established H1299 cell lines stably expressing sh-RBM15 or sh-NC-RBM15, with knockdown efficiency verified by qRT-PCR and western blot (Fig. 4A, B). Nude mouse xenograft assays showed that tumors derived from sh-RBM15-expressing cells were significantly smaller and weighed considerably less than tumors derived from control cells (Fig. 4C–F). H&E staining revealed that tissue architecture was preserved across groups (Fig. 4G), while immunohistochemical staining for Ki-67 demonstrated that proliferative activity was markedly reduced in the sh-RBM15 group relative to that in the control group (Fig. 4H). Together, these findings confirmed that RBM15 knockdown suppresses LUAD tumor growth *in vivo*.

**Fig 4.**
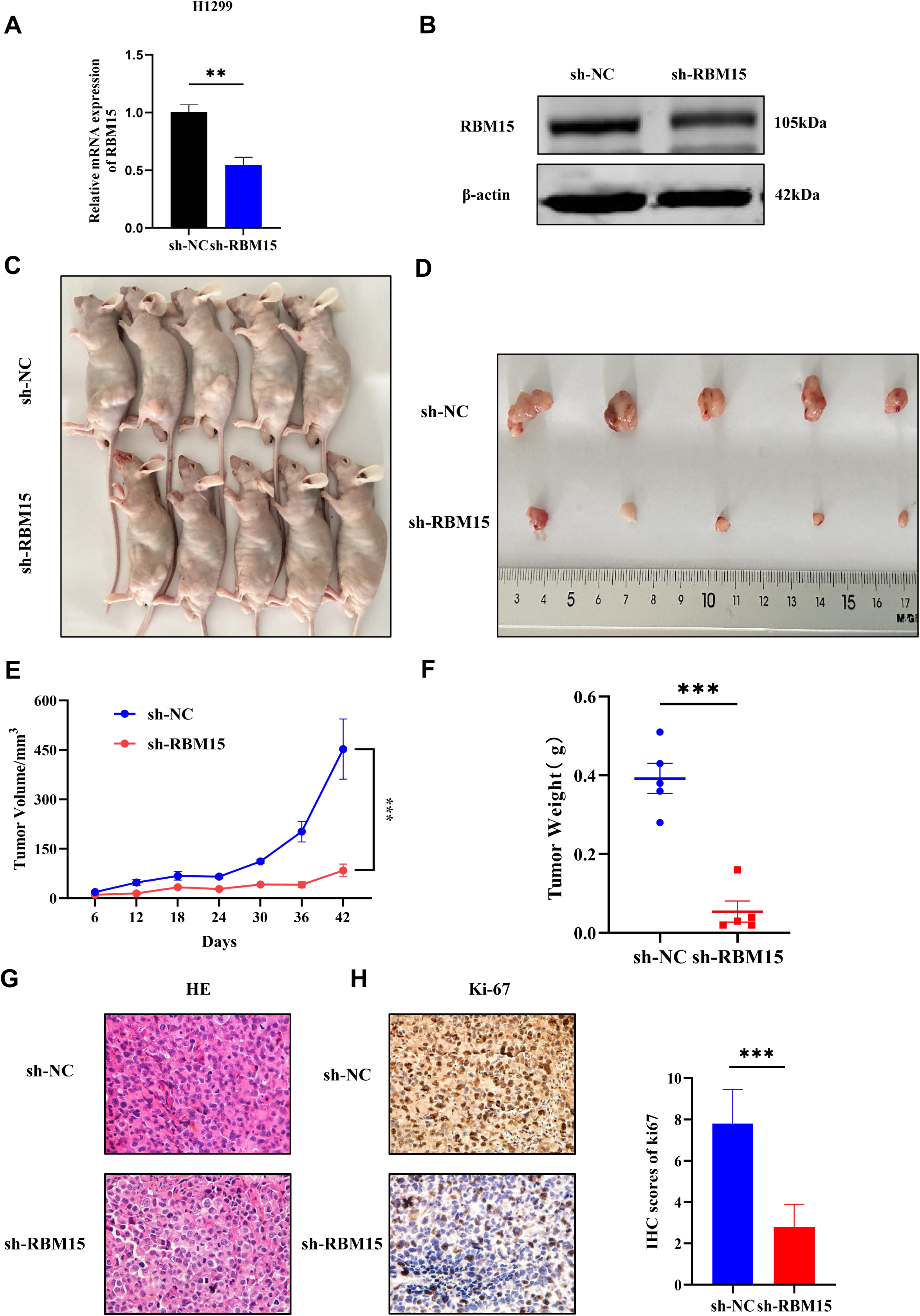
RBM15 regulates tumorigenesis and tumor growth in a nude mouse LUAD model. (A, B) mRNA and protein expression levels in stably transfected cell lines as determined by qRT-PCR and western blot. (C, D) Images of nude mice and excised tumors. (E) Tumor growth curves. (F) Tumor weights. (G) Images showing H&E-stained tumor tissue sections (×400 magnification). (H) Immunohistochemical staining for Ki-67 in tumor tissue sections (×400 magnification). \*\**P* < 0.01, \*\*\**P* <0.001.

### STIL is a target of RBM15-mediated m6A modification in LUAD

To elucidate the molecular mechanism underlying the function of RBM15 in LUAD, we undertook RNA-seq and MeRIP-seq analyses in RBM15-knockdown H1975 cells. RNA-seq results revealed that 2,964 genes were downregulated following RBM15 depletion, with 384 showing significant downregulation based on cutoff criteria of log2FoldChange < −0.5 and FDR < 0.05 (Fig. 5A, B). Meanwhile, MeRIP-seq identified 729 m6a peaks showing increased intensity and 3,933 peaks exhibiting decreased intensity in RBM15-depleted cells, indicating that RBM15 silencing markedly decreased global m6A methylation levels (Fig. 5C). There was an enrichment of m6A sites at the consensus GGAC motif, and these were predominantly localized to CDSs and 3′UTRs (Fig. 5D–F). An integrative analysis of RNA-seq and MeRIP-seq data identified 2,990 genes corresponding to m6A peaks showing decreased intensity, of which 38 overlapped with the 384 downregulated transcripts (Fig. 5G). Subsequent screening using GEPIA2 and GeneCards identified nine candidate genes that were highly expressed in LUAD, were primarily localized to the nucleus and cytoplasm, and were negatively correlated with overall survival. Visual analysis using IGV indicated that the m6A mark was highly enriched on the transcripts of four genes—*STIL*, *H2AFX*, *ZNF780A*, and *ZCCHC10* (Fig. 5H). Functional validation of these four genes in A549 and H1299 cells demonstrated that only *STIL* expression was significantly altered in response to RBM15 modulation, both overexpression and knockdown (Fig. 5I). Collectively, these findings indicated that *STIL* is a key target gene regulated by RBM15-mediated m6A modification in LUAD.

**Fig 5.**
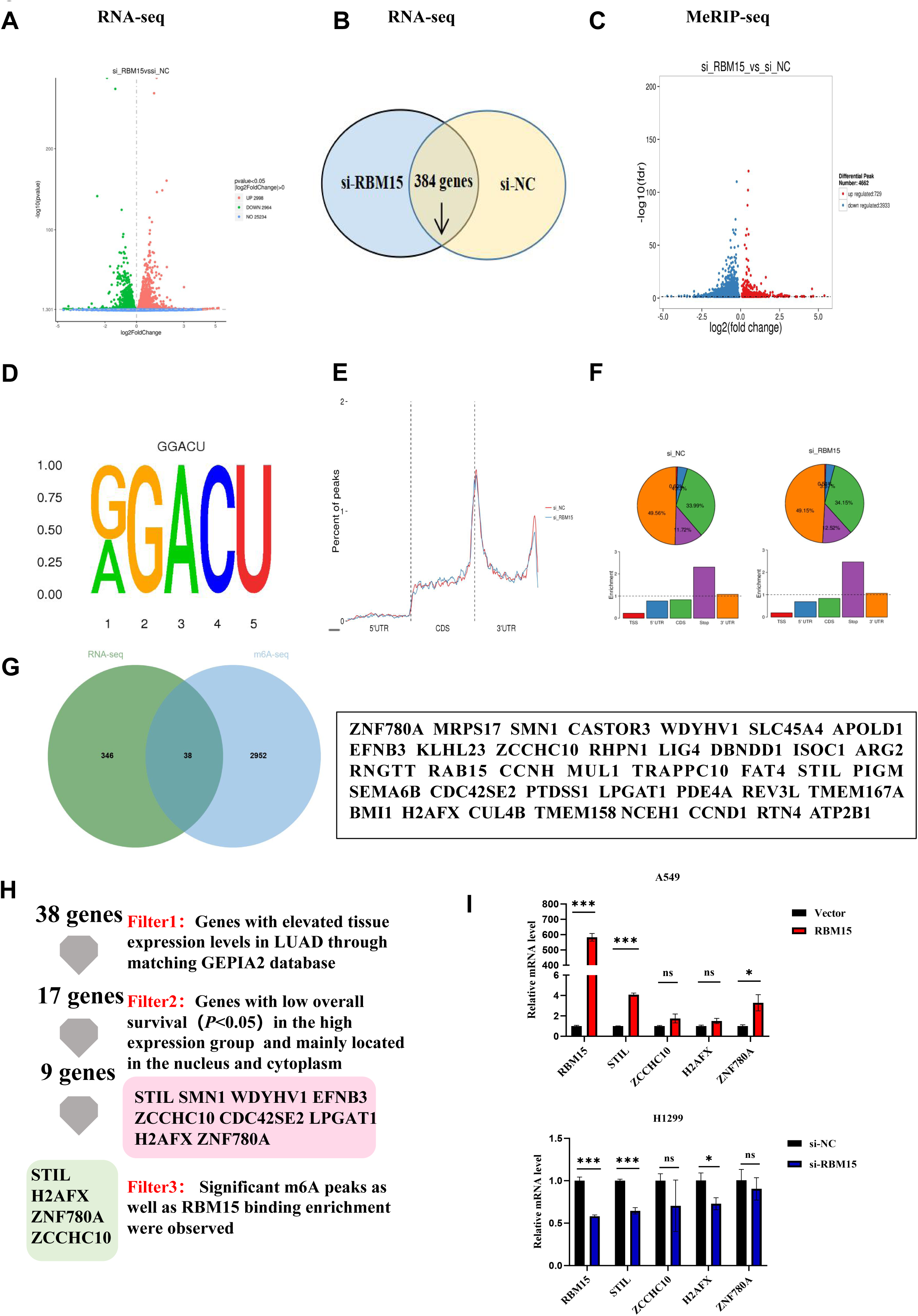
*STIL* is a target gene of RBM15. (A, B) RNA-seq analysis showing gene expression changes following RBM15 knockdown. A total of 384 genes were found to be downregulated. (C) MeRIP-seq results showing that the number of m6A peaks was significantly reduced in the si-RBM15 group compared with that in the si-NC-RBM15 group. (D–F) MeRIP results indicated that m6A marks were highly abundant in GGAC sequences, and were primarily located in coding sequences (CDSs) and 3′UTR regions of mRNA. (G) Integrated RNA-seq and MeRIP-seq analysis identified 38 candidate target genes. (H) A flowchart of the screening process leading to the identification of four candidate RBM15 target genes in H1975 cells. (I) qRT-PCR analysis showing the changes in the mRNA levels of the four candidate target genes after RBM15 overexpression and knockdown. \**P* < 0.05, *ns* indicates no significant difference.

### RBM15 regulates STIL expression *via* an m6A-dependent mechanism

Analysis of public datasets confirmed that STIL expression is significantly elevated in LUAD tissues and is negatively correlated with patient survival (Fig. 6A, C). STIL was predicted to localize to both the nucleus and cytoplasm (Fig. 6B), and its expression was markedly higher in LUAD cells than in normal lung epithelial cells (Supplementary Fig. 2A). Importantly, RBM15 knockdown reduced m6A enrichment on *STIL* transcripts (Fig. 6D). To verify whether RBM15 regulates STIL expression through m6A modification, we generated luciferase reporter constructs containing either the WT STIL sequence or a mutated (Mut) version, in which the adenine constituting the top predicted m6A site (position 2233) was replaced with cytosine (Fig. 6E). A dual-luciferase reporter assay demonstrated that RBM15 overexpression significantly enhanced luciferase activity driven by WT but not Mut *STIL* (Fig. 6F), whereas RBM15 knockdown decreased WT activity without affecting mutant sequence-driven luciferase activity (Fig. 6G). These results confirmed that RBM15 regulates STIL expression in an m6A-dependent manner. Consistent with these findings, RBM15 overexpression led to the upregulation of STIL mRNA and protein levels in A549 cells, while its knockdown resulted in the downregulation of STIL expression in H1299 cells (Fig. 6H–K). These results demonstrated that RBM15 directly regulates STIL expression through m6A-dependent methylation in LUAD.

**Fig 6.**
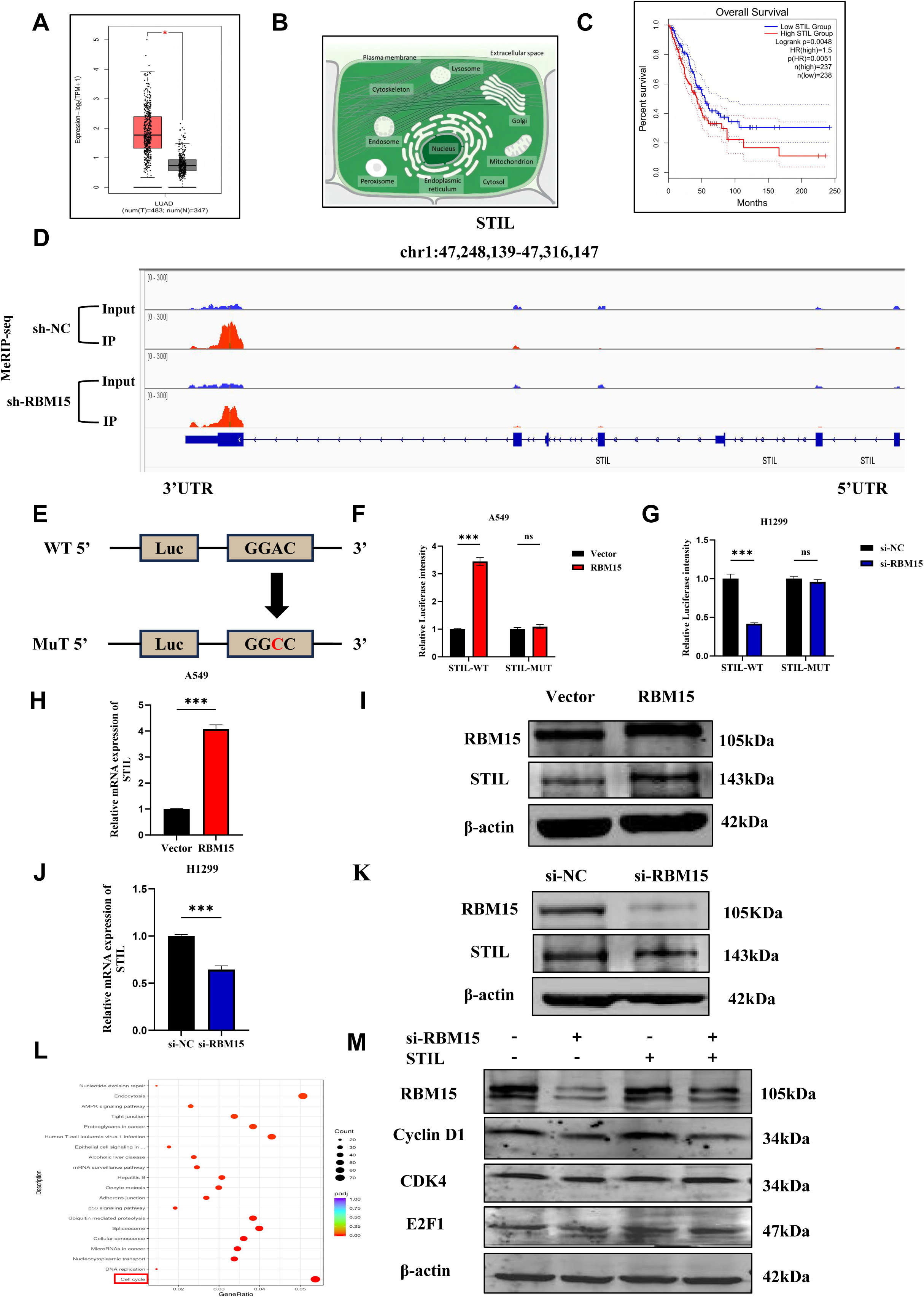
RBM15 regulates STIL expression in LUAD in an m6A-dependent manner. (A) STIL expression in LUAD based on GEPIA2 analysis. (B) STIL subcellular localization according to GeneCards. (C) The relationship between STIL expression levels and overall survival in patients with LUAD based on GEPIA2 data. (D) The downregulation of RBM15 expression reduces STIL mRNA m6A modification levels in H1975 cells. (E) Schematic illustration of luciferase reporter constructs containing wild-type STIL or its m6A-site mutant (position 2233). (F, G) Relative luciferase activity in A549 and H1299 cells under the indicated treatments. (H, J) STIL mRNA expression levels after RBM15 overexpression and knockdown. (I, K) STIL protein expression levels after RBM15 overexpression and knockdown. (L) KEGG enrichment analysis results. (M) The expression levels of cell cycle-related proteins in H1299 cells co-transfected with RBM15 siRNA and a STIL overexpression plasmid. \**P* < 0.05, \*\**P* < 0.01, \*\*\**P* < 0.001, *ns* indicates no significant difference.

### The tumorigenic effect of RBM15 in LUAD depends on STIL

KEGG pathway enrichment analysis of RBM15-regulated genes revealed that they were significantly associated with the cell cycle pathway (Fig. 6L). To further investigate the role of STIL in this process, four siRNAs targeting STIL were transfected into H1299 cells, and si-STIL (3261), which achieved the most efficient knockdown, was selected for subsequent experiments (Supplementary Fig. 2B). To determine whether STIL mediates the cell cycle-related effects of RBM15, we examined the expression of key cell cycle regulators following RBM15 silencing. In H1299 cells, RBM15 knockdown markedly reduced cyclin D1, CDK4, and E2F1 protein levels, while STIL overexpression reversed these changes (Fig. 6M). Co-transfection experiments further confirmed this STIL-mediated regulation: in A549 cells, STIL knockdown abrogated the RBM15 overexpression-induced increases in mRNA and protein levels, while in H1299 cells, STIL overexpression rescued the suppression resulting from RBM15 depletion (Supplementary Fig. 2C, D). Functionally, CCK-8 and colony formation assays showed that STIL silencing attenuated RBM15-induced proliferation in A549 cells, while STIL overexpression in H1299 cells reversed these RBM15 knockdown-induced effects (Fig. 7A, B, F, G). Similarly, wound-healing and Transwell assays demonstrated that STIL depletion diminished the enhanced migratory and invasive abilities conferred by RBM15 overexpression, whereas STIL overexpression rescued the loss of migratory and invasive capacities observed in RBM15-silenced cells (Fig. 7C–E, H–J). Collectively, these results indicated that STIL mediates the oncogenic effects of RBM15 and is required for the RBM15-mediated regulation of LUAD cell proliferation, migration, and invasion.

**Fig 7.**
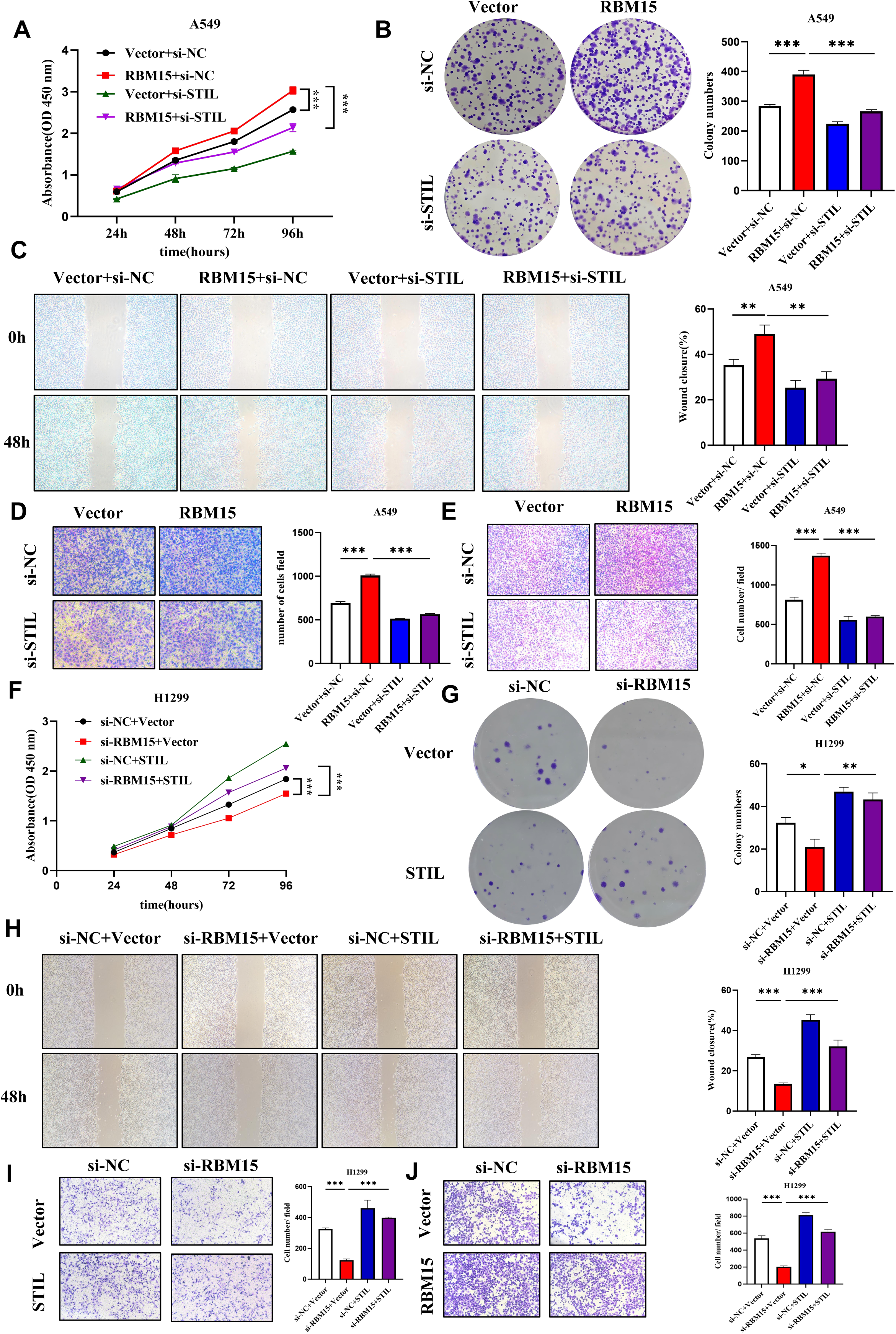
The carcinogenic activity of RBM15 is dependent on STIL. (A, B) CCK-8 and colony formation assays evaluating the effects of STIL knockdown on RBM15 overexpression-induced cell proliferation. (C, D) Wound healing and Transwell migration assays assessing the effects of STIL knockdown on RBM15 overexpression-induced cell migration. (E) Transwell invasion assays examining the effects of STIL knockdown on the RBM15 overexpression-induced increase in the invasive capacity of LUAD cells. (F–J) The effects of STIL on H1299 cells following RBM15 overexpression or knockdown were assessed using CCK-8 assays (F), colony formation assays (G), wound healing assays (H), Transwell migration assays (I), and Transwell invasion assays (J). \**P* < 0.05, \*\**P* < 0.01, \*\*\**P* < 0.001.

### RBM15 promotes LUAD progression through influencing cell cycle progression

To further confirm that RBM15 exerts its biological function *via* its effects on the cell cycle, we treated H1299 cells with the CDK4/6 inhibitor palbociclib and then divided the cells into four groups: si-NC-RBM15+DMSO, si-RBM15+DMSO, si-NC-RBM15+palbociclib, and si-RBM15+palbociclib. The IC_50_ value for palbociclib was first determined by CCK-8 assay (Fig. 8A). Subsequent proliferation assays showed that RBM15 knockdown potentiated the inhibitory effect of palbociclib on LUAD cell growth (Fig. 8B, C). Additionally, flow cytometric analysis revealed that the silencing of RBM15 led to cell cycle arrest at the G2/M phase, whereas palbociclib treatment caused arrest at the G0/G1 phase, consistent with the effect observed following CDK4/6 inhibition (Fig. 8D). Combined treatment (si-RBM15+palbociclib group) further suppressed LUAD cell proliferation, suggesting that a synergistic inhibitory effect was achieved. These findings demonstrated that RBM15 promotes LUAD progression primarily through the activation of the cyclin D1/CDK4 axis, and that pharmacologic CDK4/6 inhibition effectively blocks this pathway.

**Fig 8.**
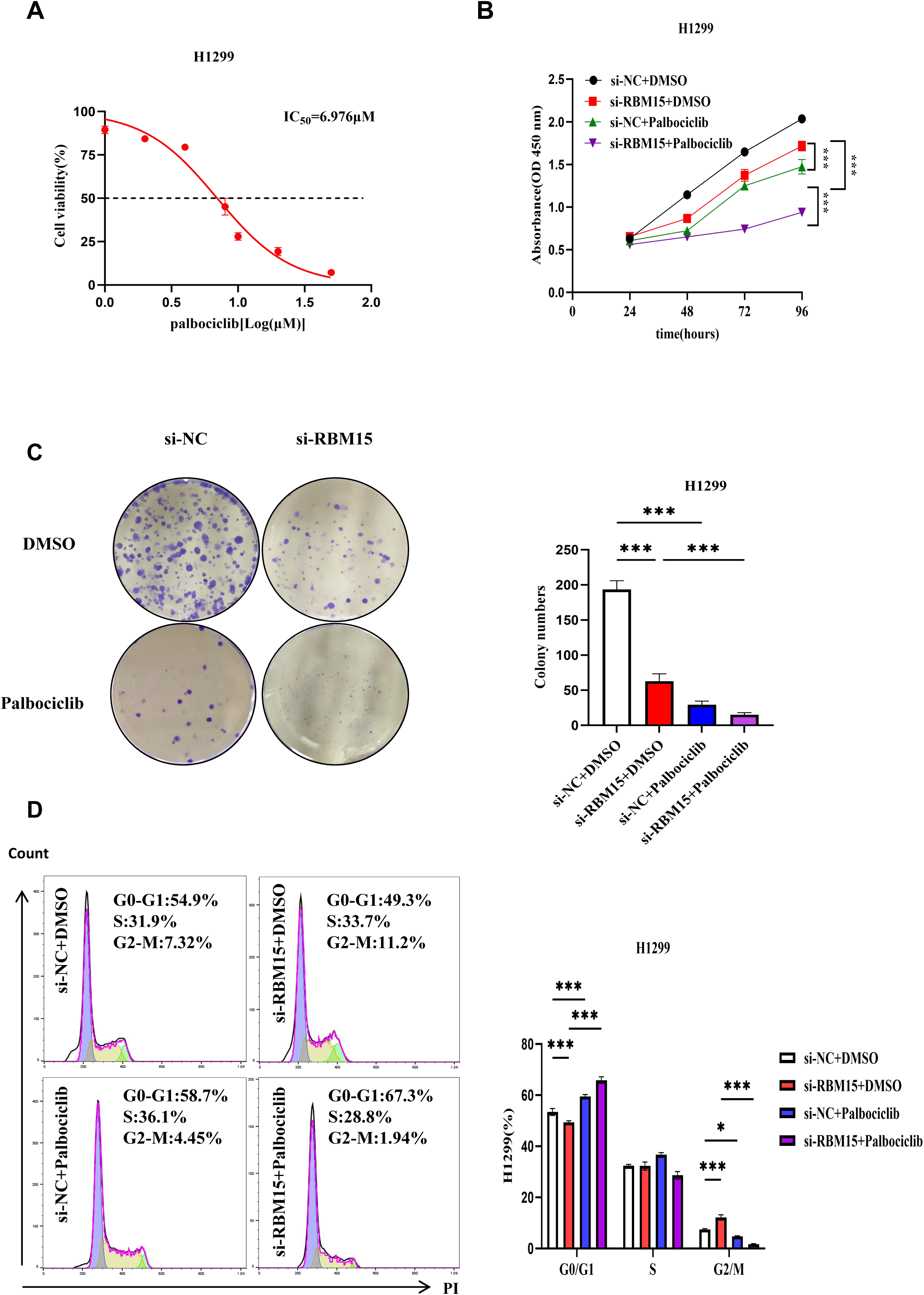
The knockdown of RBM15 enhances the inhibitory effect of palbociclib on the proliferation of LUAD cells and increases their sensitivity to the drug. (A) The IC_50_ values of palbociclib in H1299 cells were determined using the CCK-8 assay after treatment with varying drug concentrations. (B) CCK-8 assays assessing the proliferative capacity of H1299 cells in the different treatment groups. (C) Colony formation assays evaluating the proliferative capacity of H1299 cells in the different treatment groups. (D) Flow cytometric analysis of the cell cycle distribution in H1299 cells treated with different concentrations of palbociclib. \**P* < 0.05, \*\*\**P* < 0.001.

## Disscusion

Lung cancer remains the leading cause of cancer-related mortality worldwide[9]. Among its pathological subtypes, NSCLC accounts for approximately 85% of all lung cancer cases, representing a major clinical and social burden. In China, LUAD has become the predominant subtype, exhibiting consistently rising incidence and mortality rates. Prognosis and treatment strategies for NSCLC largely depend on the stage at diagnosis. While early-stage disease can often be cured by surgery, most patients are diagnosed at advanced or metastatic stages, rendering surgical intervention ineffective. Owing to the lack of highly sensitive and specific biomarkers, many patients miss the optimal therapeutic window and present with widespread metastasis, severely compromising quality of life and prognosis while imposing significant economic and psychological burdens[10]. Although current therapeutic strategies, including radiotherapy, chemotherapy, and immunotherapy, have improved survival, issues such as recurrence, toxicity, and drug resistance persist. Therefore, there is an urgent need to identify novel molecular targets and elucidate their regulatory mechanisms to improve early diagnosis and develop more effective therapeutic strategies for LUAD.

With the advancement of molecular oncology, small-molecule inhibitors targeting key signaling pathways have become an important component of cancer therapy. Many multi-targeted kinase inhibitors have shown substantial clinical efficacy and have been approved for use in the treatment of advanced, recurrent, or metastatic malignancies[11]. Continued improvements in therapeutic outcomes require a deeper understanding of the molecular mechanisms underlying tumor initiation and progression. In this study, we identified RBM15 as a key oncogenic factor in LUAD. RBM15 expression was significantly elevated in LUAD tissues and cell lines, and its expression correlated with advanced tumor stage and poor differentiation. Both i*n vitro* and *in vivo* functional assays demonstrated that RBM15 promotes LUAD cell proliferation, migration, and invasion, while concomitantly suppressing apoptosis. Mechanistically, we identified STIL as a direct target of RBM15 and found that it mediates the oncogenic effects of RBM15 through the cyclin D1/CDK4 signaling pathway. Furthermore, the pharmacologic inhibition of this pathway with palbociclib attenuated RBM15-mediated tumorigenic activity, suggesting that RBM15 may serve as a promising therapeutic target in LUAD.

M6A is the most prevalent internal RNA modification in eukaryotic mRNA and plays a pivotal role in RNA metabolism, including splicing, degradation, export, and translation[12]. This modification is dynamically regulated by methyltransferases (“writers”), demethylases (“erasers”), and binding proteins (“readers”), and dysregulation of these components has been closely linked to tumorigenesis[13,14]. For instance, elevated expression of the m6A demethylase FTO was shown to promote leukemogenesis, whereas its pharmacological inhibition elicited the opposite effect[15]. In NSCLC, METTL3 was demonstrated to enhance the translation of the oncogene *BRD4* through m6A modification and mRNA looping with EIF3h, thereby facilitating ribosomal recycling and tumor progression[16]. Similarly, WTAP has been reported to promote gastric cancer progression by upregulating MAP2K6 expression through m6A-dependent mechanisms[17]. These findings collectively underscore the critical role of m6A modification in cancer development.

RBM15, a component of the m6A methyltransferase complex, catalyzes m6A deposition in concert with METTL3, METTL14, and WTAP. Studies have shown that RBM15 plays an oncogenic role in multiple malignancies. For example, RBM15 promotes malignant progression in laryngeal squamous cell carcinoma by increasing m6A modification levels in TMBIM6 mRNA[6]. In cervical cancer, meanwhile, RBM15 knockdown decreases OTUB2 m6A methylation, leading to the inactivation of the AKT/mTOR pathway and reduced cell proliferation[18]. Consistent with these reports, our study demonstrated that RBM15 is markedly upregulated in LUAD and promotes tumor progression *via* the modulation of cell proliferation, migration, and invasion, along with cell cycle transition. Importantly, bioinformatics and sequencing analyses identified STIL as a potential downstream target of RBM15-mediated m6A modification.

STIL is a key regulator of centriole duplication and plays an essential role in the maintenance of genomic stability. Dysregulated STIL expression has been linked to aberrant cell cycle progression and oncogenic transformation. Here, we demonstrated that RBM15 enhances STIL mRNA stability through m6A modification, thereby activating the cyclin D1/CDK4 pathway and driving LUAD cell proliferation. This finding aligns with previous studies showing that STIL regulates cell cycle progression by modulating centrosome number[19]. Moreover, STIL promotes tumor growth through the IGF-1/PI3K/AKT pathway in gastric cancer and the PI3K/AKT/mTOR pathway in bladder cancers, suggesting that its oncogenic function is context-dependent[20,21]. Our rescue experiments confirmed that STIL mediates the oncogenic effects of RBM15 on the LUAD cell cycle, establishing the RBM15-STIL-cyclin D1/CDK4 axis as a key regulatory pathway in LUAD.

Additionally, treatment with the CDK4/6 inhibitor palbociclib significantly inhibited LUAD cell proliferation, further supporting the functional relevance of this pathway in this lung cancer subtype. Given the reversibility of the m6A modification, targeting m6A regulators offers a promising therapeutic avenue for LUAD. For example, METTL3 inhibition was shown to suppress leukemia progression[22], while FTO modulation enhanced anti-PD-1 efficacy in solid tumors[23]. Similarly, CDK4/6 inhibitors, such as palbociclib, have demonstrated therapeutic potential across multiple tumor types[24,25].

Despite our promising findings, several limitations of this study should be acknowledged. The sample size in this study was limited, and validation *via* additional *in vivo* and multi-center clinical studies is needed to strengthen our conclusions. Nevertheless, our results revealed a novel mechanistic insight into LUAD pathogenesis: RBM15 drives LUAD progression through the m6A modification-dependent stabilization of STIL mRNA and subsequent activation of the cyclin D1/CDK4 pathway. Targeting this regulatory axis may represent a potential strategy for improving prognosis and treatment outcomes for LUAD patients.

## Materials and methods

### Cases and specimens

This study included 43 patients with LUAD who underwent surgical resection at the Department of Thoracic Surgery, First Hospital of Hebei Medical University, between 2017 and 2023. The study materials consisted of LUAD tissues and matched adjacent normal tissues. All LUAD tissues underwent postoperative pathological diagnosis. None of the patients had received radiotherapy, chemotherapy, or biological therapy before surgery. All the patients provided written informed consent. The study was approved by the Ethics Committee of the First Hospital of Hebei Medical University (Ethics Approval Number: S01089).

### Immunohistochemistry (IHC)

Conventional paraffin sections were prepared from human tissue samples and were submitted to immunohistochemical analysis using a rabbit/mouse two-step immunohistochemistry kit (Beijing Zhongsheng Biotechnology Co., Ltd, Beijing, China). The primary antibodies used were anti-RBM15 (dilution 1:200; Proteintech, Wuhan, Hubei, China; 10587-1-AP) and anti-Ki-67 (dilution 1:50; Proteintech; 27309-1-AP). All results were evaluated using a double-blind method. Staining intensity and extent were determined based on a comprehensive scoring system.

### Cell culture and drugs

The human normal lung epithelial cell line BEAS-2B and the human LUAD cell lines NCI-H1975, NCI-H1299, and A549 were purchased from Wuhan Procell Biotechnology Co., Ltd, and validated by short tandem repeat (STR) analysis. BEAS-2B cells were cultured using DMEM (Gibco, Gaithersburg, MD, USA), and the three LUAD cell types were cultured using RPMI1640 medium (Gibco). Both media were supplemented with 10% fetal bovine serum (FBS; Gibco) and 1% penicillin-streptomycin (Solarbio Sciences & Technology Co., Ltd, Beijing, China). All cells were cultured in an incubator at 37°C with 5% CO_2_. The CDK4/6 inhibitor, palbociclib, was purchased from MedChemExpress (MCE).

### Establishment of stably transfected cell lines

Cells were transfected at 50%–70% confluence using Lipofectamine 3000 (Invitrogen, Carlsbad, CA, USA). H1299 and H1975 cells were transfected with siRNA-RBM15 (si-RBM15) or siRNA negative control (si-NC-RBM15) (GenePharma Co., Ltd, Shanghai, China), while A549 and H1975 cells were transfected with the pcDNA3.1-RBM15 plasmid (RBM15) or the negative (vector only) control. A549 cells were transfected with siRNA-STIL or siRNA negative control (si-NC-STIL) (GenePharma), while H1299 cells were transfected with the pcDNA3.1-STIL plasmid (STIL) or the negative (vector only) control (GenePharma). After transfection, cells were cultured for 48 h for subsequent experiments. H1299 cells were transfected with RBM15 shRNA (sh-RBM15) or shRNA negative control (sh-NC) (GenePharma), followed by cell selection using 400 μg/mL G418.

### RNA extraction and quantitative reverse transcription-PCR (qRT-PCR)

Total RNA was extracted from cells and tissues using RNA-Easy Isolation Reagent (Vazyme Biotechnology Co., Ltd, Nanjing, China), and then reverse-transcribed into cDNA using the PrimeScript RT Kit (Takara, Beijing, China) according to the manufacturer’s instructions. qPCR was performed using AceQ Universal SYBR qPCR Master Mix (Vazyme), with the β-actin gene serving as the internal reference. The relative expression levels of each sample were calculated using the 2-^ΔΔCT^ method. The primers used for qPCR are listed in Supplementary Table 1.

### Western blot

Total protein was extracted from cells using RIPA lysis buffer (Beyotime Biotechnology Co., Ltd, Shanghai, China). Protein (20–30 μg per sample) was separated by 10% SDS-PAGE (Bio-Rad Laboratories Inc., Hercules, CA, USA), transferred to a PVDF membrane for 40 min, blocked with 5% non-fat milk for 2 h, and incubated overnight at 4°C with primary antibodies targeting RBM15 (dilution 1:2000, 10587-1-AP), STIL (dilution 1:2000, 66876-1-Ig), E2F1 (dilution 1:250, 66515-1-Ig), cyclin D1 (dilution 1:250, 60186-1-Ig), CDK4 (dilution 1:500, 11026-1-AP) (all Proteintech), β-actin (dilution 1:1000, ZSBG-Bio, TA-09). The PVDF membrane was then incubated with goat anti-rabbit or goat anti-mouse fluorescent secondary antibodies, and the immunoreactive bands were detected using the Odyssey scanning system (LICOR Biosciences, Lincoln, NE, USA).

### Cell proliferation assay

Cells from each group were seeded at a density of 2 × 10^3^ cells per well in a 96-well plate and cultured in complete medium in a cell culture incubator. After 24, 48, 72, and 96 h, 10 μL of Cell Counting Kit-8 reagent (CCK-8; Dojindo, Tokyo, Japan) was added to each well, and culture was continued for 2 h. The absorbance at 450 nm of each well was subsequently measured using a Promega GloMax fluorescence detector (Promega, Madison, WI, USA).

### Colony formation assay

Cells were seeded at a density of 500 cells per well in a six-well plate for 10 days, with the culture medium changed every 3 days. After washing two to three times with PBS, the cells were fixed in 4% paraformaldehyde for 30 min, stained with 0.1% crystal violet for 20 min, and then counted.

### Wound healing assay

In the wound healing assay, cells from each group were seeded in six-well plates. At 100% cell confluence, two straight lines were drawn in each well using a 200 μL pipette tip. After washing twice with PBS, cell culture was continued in serum-free medium, and cells were imaged at 0 h and 48 h, followed by calculation of the wound healing rate.

### Transwell migration and invasion assays

Cell migration and invasion assays were performed using Transwell chambers (Corning Incorporated, Corning, NY, USA). In the migration assay, 4 × 10^4^ cells were suspended in 200 μL of serum-free RPMI 1640 medium and transferred to the upper chamber, while 600 μL of complete medium was placed in the lower chamber to induce chemotaxis. In the invasion assay, the upper chamber contained 100 μL of medium plus cells, while the lower chamber contained 600 μL of medium; other conditions were the same as in the cell migration assay. After incubation for 24 h, the cells were fixed in 4% paraformaldehyde for 30 min, stained with 0.1% crystal violet for 20 min, imaged, and counted using ImageJ software.

### Apoptosis detection

To quantify apoptosis, 48 h after transfection, both the supernatant and adherent cells were collected from each group, washed two/three times with PBS, and then stained using the Annexin V-FITC/PI apoptosis detection kit from NeoBioscience (Shenzhen, China). After 15 min, the cells were analyzed using a flow cytometer (BD Biosciences, San Jose, CA, USA). Finally, the apoptosis rate was determined using FlowJo software.

### Cell cycle analysis

The preliminary steps for cell cycle analysis were the same as those used for apoptosis analysis. Cells from each group were processed using the Cell Cycle and Apoptosis Analysis Kit (Beyotime, C1052). After collection, the cells were fixed in 70% ethanol and incubated at 4°C for 2 h. Subsequently, the cells were stained with 0.5 mL of propidium iodide in the dark at a constant temperature for 30 min. Finally, red fluorescence was detected using a flow cytometer at an excitation wavelength of 488 nm, and cell cycle profiles were analyzed using FlowJo software.

### RNA sequencing and methylated RNA immunoprecipitation (MeRIP) sequencing analysis

Total RNA was extracted from H1975 cells previously transfected with si-RBM15 or si-NC-RBM15. The RNA samples were sent to Novogene for sequencing, which was performed on an Illumina sequencing platform, yielding raw data. For MeRIP-seq, RNA libraries were constructed and subsequently sequenced at Novogene. The procedure involved fragmenting the mRNA and immunoprecipitating m6A-modified RNA fragments using antibodies specific for the m6A modification. The immunoprecipitated RNA fragments were subsequently subjected to high-throughput sequencing. Statistical analysis was conducted on the peak distribution of mRNA transcripts across the 5′UTR, coding sequence (CDS), 3′UTR, and other functional regions.

### Animal studies

Ten 5-week-old SPF-grade BALB/c nude mice were purchased from Beijing Huafukang Biotechnology Co., Ltd. The animals were randomly divided into a negative control (sh-NC-RBM15) group and a *RBM15* gene knockdown (sh-RBM15) group. H1299 cells (2 × 10^6^) stably transfected with sh-NC-RBM15 or sh-RBM15 were subcutaneously injected into the left hind limb of nude mice. Tumor size was measured every 6 days starting from day 4 post-injection. When the tumor volume approached 1000 mm^3^, mice were humanely euthanized. Tumor tissues were collected, weighed, fixed in formalin, embedded in paraffin, sectioned, and subjected to hematoxylin and eosin (H&E) staining and immunohistochemical analysis. All animal-related protocols were approved by the Ethics Committee of the First Hospital of Hebei Medical University (Ethics Approval Number: S01090).

### Luciferase reporter gene assay

Primers were designed to generate fragments corresponding to wild-type (WT) and mutant (Mut) target sequences within the *STIL* gene. Double-stranded DNA fragments were generated *via* annealing, ligated into a linearized pmirGLO vector, and transformed into XL1-Blue recipient cells. Positive clones were screened and sequenced for validation. H1299 and A549 cells were seeded in 96-well plates and cultured to 70%–80% confluence. The cells were then co-transfected with plasmids containing either WT or Mut STIL sequences and RBM15/si-RBM15 (or controls). After 48 h of transfection, the cells were lysed, and firefly and *Renilla* luciferase activity was measured using a dual luciferase reporter gene assay kit (Beyotime, RG009). Relative activity (RLU1/RLU2 ratio) was detected using a GloMax 96 fluorescence detector (Promega, USA).

### Bioinformatics analysis

RBM15 mRNA expression levels in LUAD tissues and adjacent normal tissues were analyzed using public data from The Cancer Genome Atlas (TCGA) (https://www.aclbi.com/static/index.html#/tcga) and GeneExpression Omnibus (GEO) (https://www.aclbi.com/static/index.html#/geo) databases. The MicroBioinformatics online cloud platform (http://www.bioinformatics.com.cn/) was employed for GO functional and KEGG pathway enrichment analyses of preliminarily screened candidate genes. PrimerBank (https://pga.mgh.harvard.edu/primerbank/) and Primer Design Tool (https://www.ncbi.nlm.nih.gov/tools/primer-blast/) were used for primer prediction. GEPIA2 (http://gepia2.cancer-pku.cn/#index) was used to investigate gene expression levels in LUAD tissues and adjacent normal tissues, as well as for survival curve analysis. Subcellular localization of proteins encoded by the genes of interest was assessed using GeneCards (https://www.genecards.org/). IGV software was used for the visual analysis of m6A methylation levels and RBM15 binding enrichment.

### Drug sensitivity

The IC_50_ value of palbociclib was first established in WT H1299 cells using a range of drug concentrations. Subsequently, transfected H1299 cells were seeded in 96-well plates for 24 h, treated with different doses of palbociclib for 2 days, and then subjected to a CCK-8 assay for determination of cell viability. For the colony formation assay, transfected H1299 cells were seeded in six-well plates and treated with different doses of palbociclib for 10 days, and then fixed and stained with crystal violet. To assess the effect of the drug on the cell cycle of LUAD cells, transfected H1299 cells were divided into different subgroups, treated with palbociclib for 2 days, and then analyzed by flow cytometry.

### Statistical analysis

Normally distributed data were expressed as means ± standard deviation, while non-normally distributed data were expressed as medians with interquartile ranges. *P*-values were determined using the *t*-test, Fisher’s exact test, and the chi-square test. Statistical analysis was performed using GraphPad Prism 10.1.2 (GraphPad Software, La Jolla, CA, USA) and SPSS 26.0 (IBM, Armonk, NY, USA). *P* < 0.05 was considered statistically significant.

## Data availability

The datasets generated during and/or analysed during the current study are available from the corresponding author on reasonable request.

## Author contributions

The experiment was conducted by YFC, JJZ, and HJY, followed by data analysis and manuscript drafting. LHW and MHL jointly participated in the study design and contributed to manuscript revisions. LW, JLM, and JW participated in the experiment. JW, TL, and WFY were responsible for study design and supervision, while also contributing to manuscript revisions.

## Supplementary Tables

**Supplementary Table 1.** The sequences of the primers used for the amplification of RBM15, STIL, ZCCHC10, H2AFX, ZNF780A, and β-actin.

**Supplementary Table 2.** Analysis of clinicopathological data.

