## Supplementary figures and images for "RBM15 promotes lung adenocarcinoma progression and palbociclib sensitivity *via* m6A-mediated STIL activation and downstream cyclin D1/CDK4 signaling"

### Supplementary Figure 1

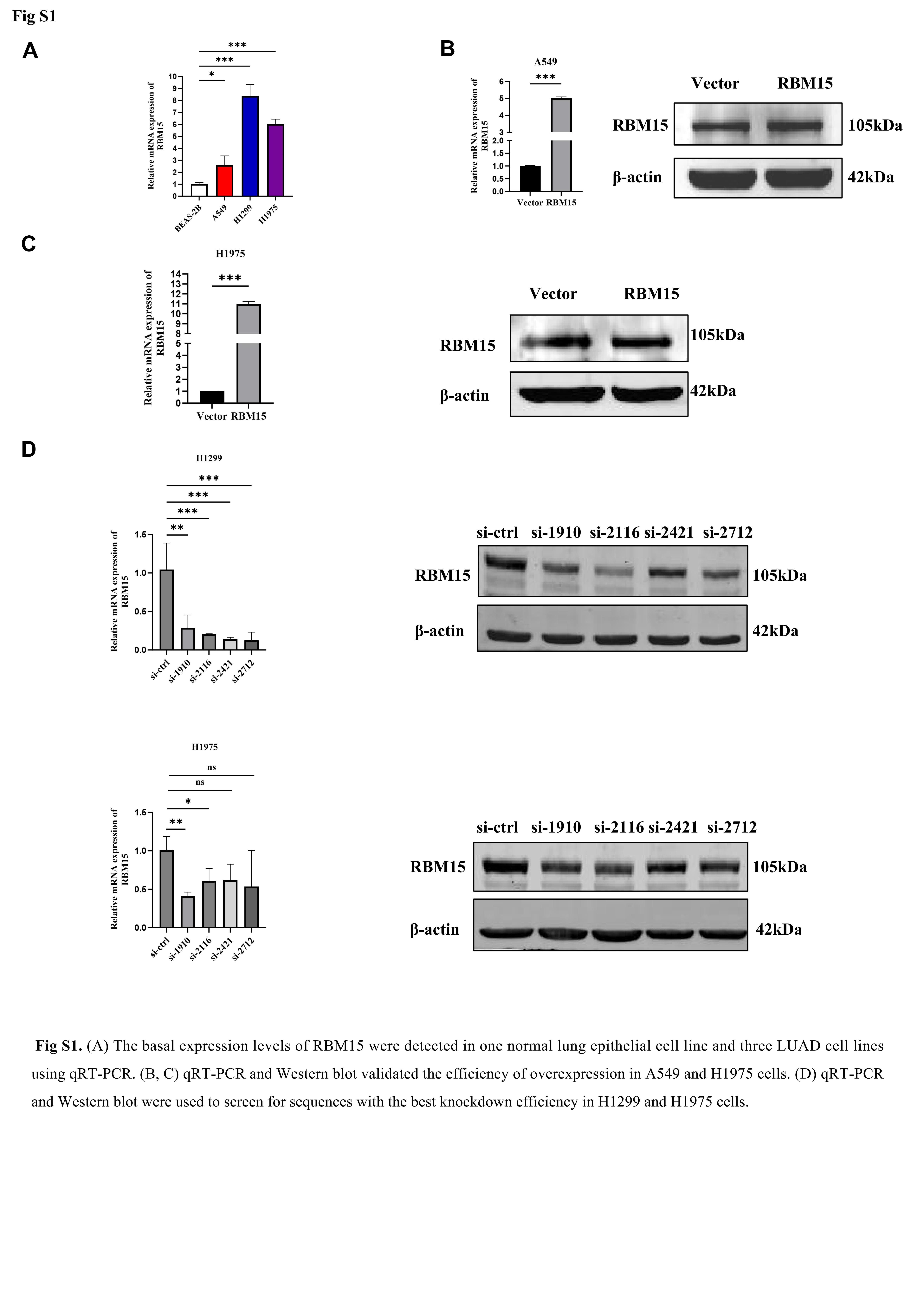

### Supplementary Figure 2

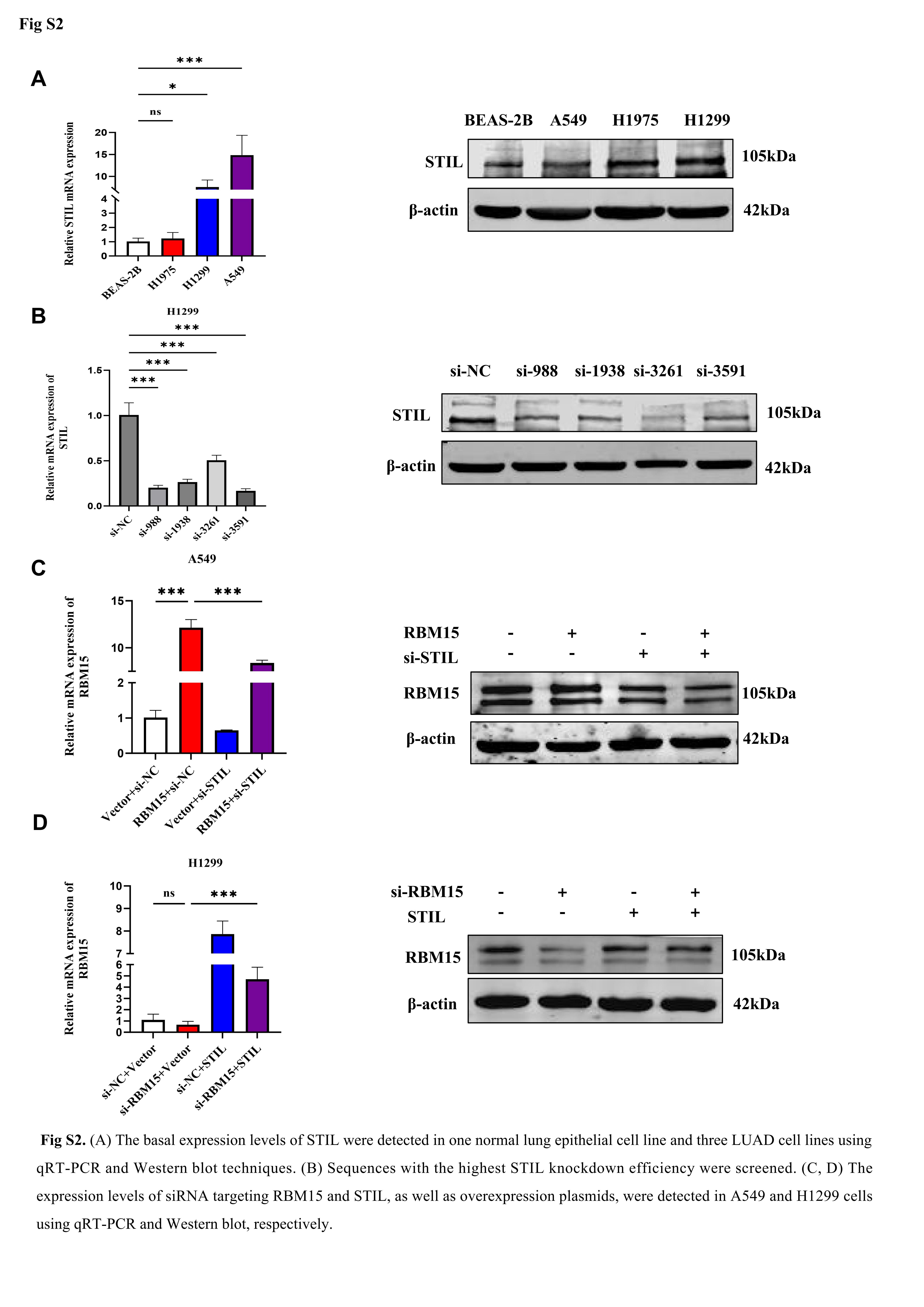
