## Supplementary Tables for "RBM15 promotes lung adenocarcinoma progression and palbociclib sensitivity *via* m6A-mediated STIL activation and downstream cyclin D1/CDK4 signaling"

**Table S1.** The sequences of the primers used for the amplification of RBM15, STIL, ZCCHC10, H2AFX, ZNF780A, and β-actin.

| Primer name | Sequence (5'-3') |
| --- | --- |
| β-actin-F | GAGTCAACGGATTTGGTCGT |
| β-actin-R | CATGGGTGGAATCATATTGGA |
| RBM15-F | CCTTCATGCCTTCCCACCTT |
| RBM15-R | ATAACAGGGTCAGCGCCAAG |
| STIL-F | TTTGACTTGCATTGGGCAGC |
| STIL-R | AATGGGGATGGGCTTCACAG |
| ZCCHC10-F | CAGTAGCAGTAGCAGTGACAGT |
| ZCCHC10-R | TGGTGGTTCATCGTCAGAGC |
| H2AFX-F | GGTGCTTAGCCCAGGACTTTC |
| H2AFX-R | GTCACTCGGGAGGAAGATGTG |
| ZNF780A-F | CGACGACATAGCGGGGTA |
| ZNF780A-R | AGCCCCAATCCGCAGAGAT |

**Table S2.** Analysis of clinicopathological data.

| Groups | Cases | *RBM15* expression | | Positive rate (%) | *χ*^2^ value | *P* value |
| --- | --- | --- | --- | --- | --- | --- |
|  |  | High | Low |  |  |  |
| Lung adenocarcinoma tissues | 43 | 28 | 15 | 65.1% | 28.667 | <0.001^***^ |
| Adjacent normal tissues | 43 | 4 | 39 | 9.3% |  |  |

****P*<0.001.

| Groups | Cases | *RBM15* expression | | *χ*^2^ value | *P* value |
| --- | --- | --- | --- | --- | --- |
|  |  | High | Low |  |  |
| Gender | | | | | |
| Male | 18 | 11 | 7 | 0.219 | 0.640 |
| Female | 25 | 17 | 8 |  |  |
| Age (years) | | | | | |
| ≤ 65 | 23 | 14 | 9 | 0.393 | 0.531 |
| > 65 | 20 | 14 | 6 |  |  |
| Tumor size (cm) | | | | | |
| < 3 | 36 | 25 | 11 | 1.824 | 0.177 |
| ≥ 3 | 7 | 3 | 4 |  |  |
| **Differentiation** | | | | | |
| **Medium+Low Differentiation** | 28 | 22 | 6 | 6.397 | 0.011^*^ |
| **High Differentiation** | 15 | 6 | 9 |  |  |
| Lymph node metastasis | | | | | |
| Absent | 40 | 25 | 15 | - | 0.541 |
| Present | 3 | 3 | 0 |  |  |
| **TNM staging** | | | | | |
| **I** | 35 | 20 | 15 | - | 0.036^*^ |
| **II+III+IV** | 8 | 8 | 0 |  |  |

**P*<0.05.
